# Spike-phase coupling patterns reveal laminar identity in primate cortex

**DOI:** 10.1101/2022.10.26.513932

**Authors:** Zachary W. Davis, Nicholas M. Dotson, Tom Franken, Lyle Muller, John Reynolds

## Abstract

The cortical column is one of the fundamental computational circuits in the brain. In order to understand the role neurons in different layers of this circuit play in cortical function it is necessary to identify the boundaries that separate the laminar compartments. While histological approaches can reveal ground truth they are not a practical means of identifying cortical layers *in vivo*. The gold standard for identifying laminar compartments in electrophysiological recordings is current-source density (CSD) analysis. However, laminar CSD analysis requires averaging across reliably evoked responses that target the input layer in cortex, which may be difficult to generate in less well studied cortical regions. Further the analysis can be susceptible to noise on individual channels resulting in errors in assigning laminar boundaries. Here, we have analyzed linear array recordings in multiple cortical areas in both the common marmoset and the rhesus macaque. We describe a pattern of laminar spike-field phase relationships that reliably identifies the transition between input and deep layers in cortical recordings from multiple cortical areas in two different non-human primate species. This measure corresponds well to estimates of the location of the input layer using CSDs, but does not require averaging or specific evoked activity. Laminar identity can be estimated rapidly with as little as a minute of ongoing data and is invariant to many experimental parameters. This method may serve to validate CSD measurements that might otherwise be unreliable or to estimate laminar boundaries when other methods are not practical.

## Introduction

Linear array electrodes have become a ubiquitous electrophysiological tool for understanding the functional roles of neural populations across the layers of the cortex, their interactions, and the computations they perform. This understanding requires reliable assignment of neurons to their respective laminar compartments. Precise localization of individual neurons can be obtained by electrolytic lesion to mark the position of an electrode, however this procedure is not practical in experiments where the same animal is used in multiple experimental sessions as is nearly always the case in experiments involving non-human primates. The gold standard for identifying laminar compartments from functional recordings is current-source density (CSD) analysis (Mitzdorf and Singer, 1978; Mitzdorf, 1985; Schroeder, 1998). The CSD represents the second spatial derivative of local field potential (LFP) activity averaged to repeated events. These events are often sensory-evoked stimulation as in flashed visual stimuli in visual cortex (Schroeder et al., 1991; Maier et al., 2010; Wang et al., 2020) or short duration sounds in auditory cortex (Lakatos et al., 2007; Happel et al., 2010; Szymanski et al., 2011) which produce feedforward activation of the canonical cortical columnar circuit via activation of the input layer(Mitzdorf and Singer, 1978; Mitzdorf, 1985; Gratiy et al., 2011). Laminar compartments can then be assigned from the timing of subsequent patterns of current sources and sinks across electrode contacts on the linear array (Mitzdorf, 1985). The earliest current sink reflects the feed-forward activation of the input layers (Mitzdorf, 1985, 1987), and the current sources above and below are used to estimate the boundaries with respect to superficial (layers 1-3) and deep (layers 5 and 6) cortical layers (Self et al., 2013). CSD analysis has been validated histologically(Mitzdorf and Singer, 1979; Schroeder et al., 1991; Takeuchi et al., 2011) and reliably captures known laminar differences in the functional properties of cortical layers, such as differences in evoked latencies across layers in response to driving input(Bode-Greuel et al., 1987; Einevoll et al., 2013; Plomp et al., 2017).

There are some limitations on the use of CSD for identifying cortical layers in electrophysiology recordings. CSD analysis has largely been limited to primary sensory areas where the types of sensory stimuli that can robustly and reliably generate the evoked responses necessary for averaging CSDs are well established. Although CSD analysis has been used in higher order cortical areas where sensory evoked responses are apparent such as visual responses in frontal cortex(Godlove et al., 2014; Bastos et al., 2018), it is less clear what to use as a triggering event in non-sensory cortical areas to reveal similar laminar patterns. Another limitation of CSD analysis is that depth estimates can only be taken as the average across the period of sensory response collection, making it difficult to track electrode drift during a recording. Further, noise in recordings due to bad channels, variability in filtering parameters, or ambiguity in the identifying source-sink pattern could potentially lead to the elimination of otherwise useful data or inaccurate laminar assignment. Without an alternative means of estimating the depth of electrode contacts, analyses of cortical circuit function are at risk of using biased definitions of laminar identity resulting in spurious conclusions about layer dependent neuronal features.

Here we present a new method for determining laminar boundaries in cortical recordings based on spike-phase coupling patterns across linear array electrodes. Previous work has shown ongoing spiking activity is strongly coupled to the phase of LFP fluctuations in the cortex(Eeckman and Freeman, 1990; Destexhe et al., 1999; Haegens et al., 2011; Dotson et al., 2014; Esghaei et al., 2018; Davis et al., 2022) and we hypothesized that, as phase shifts occur across the layers of the cortex, the preferred phase angle of this spike-LFP relationship should be influenced by laminar differences in sources and sinks. We find that there is indeed a phase reversal in spike-field coupling that reliably corresponds to the laminar boundary between input and deep layers as estimated by CSD. We therefore propose a novel methodology, called laminar phase coupling (LPC), for identifying the laminar boundary between input and deep cortical layers. This pattern reliably reversed in awake recordings from both common marmosets and rhesus macaque in multiple cortical areas including extrastriate visual and prefrontal cortex. This method can be applied to the same data used for CSD analysis, but can also be applied to any cortical linear array recording data without the need for specific sensory stimuli as with CSD analysis. Data recorded under a variety of conditions, such as during behavioral tasks, spontaneous activity during fixation, or continuous data recorded in the dark all reliably reveal the same pattern of laminar phase separation. LPC is largely invariant to filtering or referencing, and can be done on single- or multi-unit activity. The analysis does not require averaging and can be calculated online to estimate recording depth throughout the duration of an experiment. The preferred phase angle in the spike-field relationship is a simple yet effective measure for estimating contact depth in linear array recordings with respect to laminar boundaries and may help inform the study of cortical circuits in situations where CSD measurements are ambiguous or where CSD methods are impractical or ineffective.

## Results

We recorded electrophysiology data from 32-channel linear probes (Atlas Neuroengineering) inserted perpendicular to the cortical surface into cortical regions V4 in two macaque monkeys and MT or PFC in two marmoset monkeys performing head-fixed visual tasks under various experimental conditions (Figure 1a). CSD analyses were performed using stimulus-locked LFP epochs (Figure 1b). These epochs were locked to stimulus flashes in the case of V4 and MT recordings or full field flashes in the case of PFC recordings. As standard in CSD analysis, we took from the resulting evoked source-sink patterns the earliest sink (red) as the location of the input layers and assigned our reference depth relative to the input layer as the bottom of the current sink, with the boundaries to the sources (blue) above and below identifying the boundaries of the superficial and deep cortical layers (Figure 1c and d).

**Figure 1.**
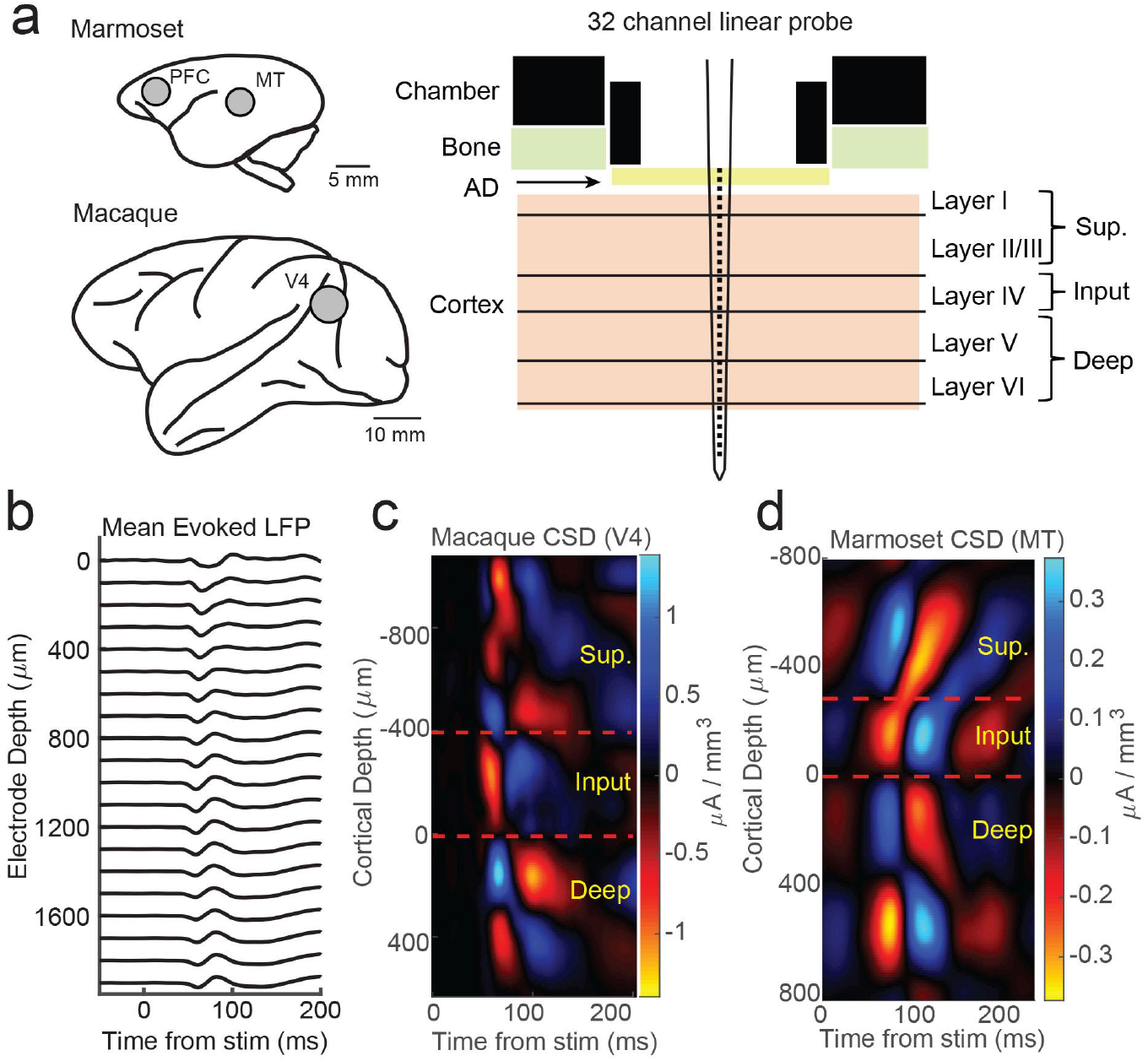
Current source density analysis reveals laminar boundaries in cortical recordings. (a). Schematic of recording locations in two marmosets (PFC or MT) and two macaques (V4). Recordings were made with a 32 channel linear silicon probe in an acute recording chamber penetrating through a silicone artificial dura (AD). (b) Average evoked LFP responses to a stimulus flash in an example macaque recording session in area V4. Each line is the response measured on a single contact. The depth is in absolute distance from the most proximal contact. (c) Current source density (CSD) measurement from the example recording session in b. The input layer is defined as the bottom and top of the earliest current sink (red), with the superficial and deep layers defined as above and below the input layer respectively. Depth is measured relative to the bottom of the input layer. (d) Same as c, but in an example marmoset MT recording session.

In order to test the relationship between spiking and LFP phase across the layers of the cortex, we first identified multi-unit spike times and measured the *generalized phase* (GP) of the LFP filtered from 5-50 Hz(Davis et al., 2020, 2022). The use of a wider filter than traditionally used (i.e. 4-8 Hz or 8-15 Hz) helps reduce phase distortions that occur when applying narrowband filters to signals that have broad spectral content. GP provides a measure of wideband phase that corrects for errors that may arise from applying the Hilbert Transform to signals with broader spectral content, although our results did not depend on our specific use of these techniques. We binned electrodes into superficial, input, and deep layers depending on their location relative to the source-sink pattern defined in the CSD analysis.

We first calculated the degree of spike coupling on each channel to the LFP phase measured on the same channel grouped by cortical layer. This was done by taking the length of the circular resultant of the spike-phase distribution, which we call the spike-phase index (SPI). This index value ranged from 0 (perfectly uniform spike-phase distribution) to 1 (all spikes occur at a single phase). The superficial channels had an average SPI value of 0.162 ± 0.012 (mean ± S. E. M.; N = 34 sessions). This was significantly greater than the input layer (SPI = 0.122 ± 0.010 mean ± S. E. M.; p = 0.0003, Wilcoxon signed-rank test), and both were significantly greater than the deep layers (SPI = 0.073 ± 0.009 S. E. M.; p = 0.000003 S. v. D. and p = 0.000009 I. v. D.). The preferred LFP phase angle (i.e. the circular mean of the phase distribution) for spiking in the superficial and input layers was −2.86 rad and −2.68 rad respectively which corresponds towards the trough of ongoing LFP fluctuations. This was significantly different from the preferred phase angle for deep layer spiking (−1.44 rad; F = 38.52, p < 1 × 10^−7^ and F = 21.15, p = 0.00002 respectively; Watson-Williams test) suggesting that indeed the deeper layers may be distinguished by a change in the preferred spike-LFP phase angle relative to the superficial electrodes and this could be read out from the spike-LFP relationship across the depth of the cortex.

While the previous analysis examined the spike-phase relationship on each channel as a function of cortical depth on average, because of variability in the preferred mean phase on each channel (due to factors such as the strength of spike-LFP coupling and the number of spikes recorded) there was no significant correlation between the preferred phase angle and the depth of the electrode as measured with CSD (circular-linear correlation r = 0.28; p = 0.24). A more sensitive measure is to relate the preferred phase angle in the spike-LFP relationship across the depths of all channels in the cortex. This was achieved by calculating the SPI from the multi-unit spiking activity on a given channel relative to the LFP phase angle measured on each contact across the depth of the recording (Figure 3a). We call the resulting matrix the cross-channel LPC. As an example, the first matrix row is derived from the spikes recorded on the first channel compared against the phase measured from the LFP on each of the 32 channels. The second row is derived from the spikes on the second channel compared against the phase measured from the LFP on each channel, and so on. The result is a 32-by-32 matrix where each cell represents the relation between the spikes on one channel and the LFP phase on another channel. However, not all of these channels are in the cortex, so we realigned the matrix based on estimates of cortical depth from the bottom of the input layer as identified from CSD analysis.

We first looked at the SPI values across the cross-channel LPC matrix. Figure 3b shows an example from a single recording (note that the diagonal in this matrix, from top left to bottom right, represents the approach in Figure 2). In this recording, we found that superficial spiking activity (defined based on CSD analysis) was strongly coupled to the LFP phase recorded on other superficial channels, which dropped off sharply below the input layer, and recovered when computed relative to the LFP phase on deeper channels. We next looked at the preferred phase angle for spiking activity on each channel relative to the LFP phase measured for different laminar compartments (Figure 3c). We found that spiking activity on all channels, regardless of cortical depth, tended to spike during ±π radian LFP phase angles measured on superficial channels. This phase relationship flipped such that spiking on all channels, regardless of cortical depth, tended to spike during 0 radian phase angles for LFP measured on deep channels. This phase reversal occurred about the channel we identified from CSD analysis as the boundary between the input and deep layers in the cortical column.

**Figure 2.**
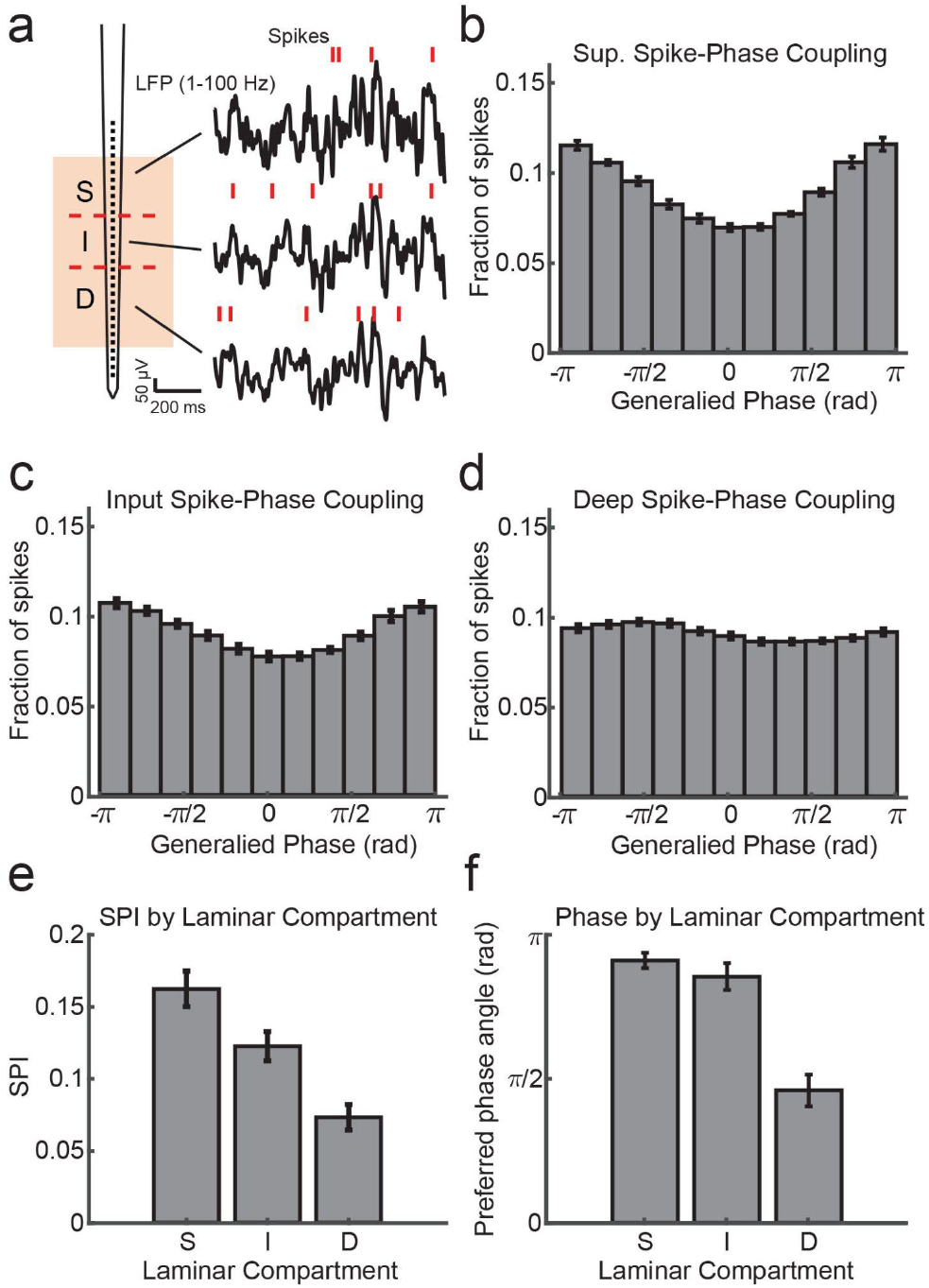
Within channel Spike-LFP phase coupling strength and preferred phase angle varies across layers. (a). Spike-LFP phase coupling was measured by taking the phase of the LFP on a contact at the times when multi-unit spikes were detected on the same contact. (b). Spike-phase distribution for superficial contacts averaged across all recordings sessions (N = 34 sessions across 4 monkeys). The spike-phase distribution was strongly peaked towards ±π rad. (c, d). Same as b, but for contacts in the input and deep layers. (e) The average spike-phase index (SPI) was significantly weaker for input and deep layers relative to the superficial layer (0.16 v. 0.12 and 0.07; p = 0.017 S. v. I. and p < 1 × 10^−7^ S. v D.; Wilcoxon signed-rank test). (f). There was no difference in the preferred LFP phase angle between superficial and input layers, but a significant difference between these layers and deep layers (−2.86 rad and −2.68 rad v. −1.44 rad; F = 38.52, p < 1 × 10^−7^ and F = 21.15, p = 0.00002 respectively; Watson-Williams test).

**Figure 3.**
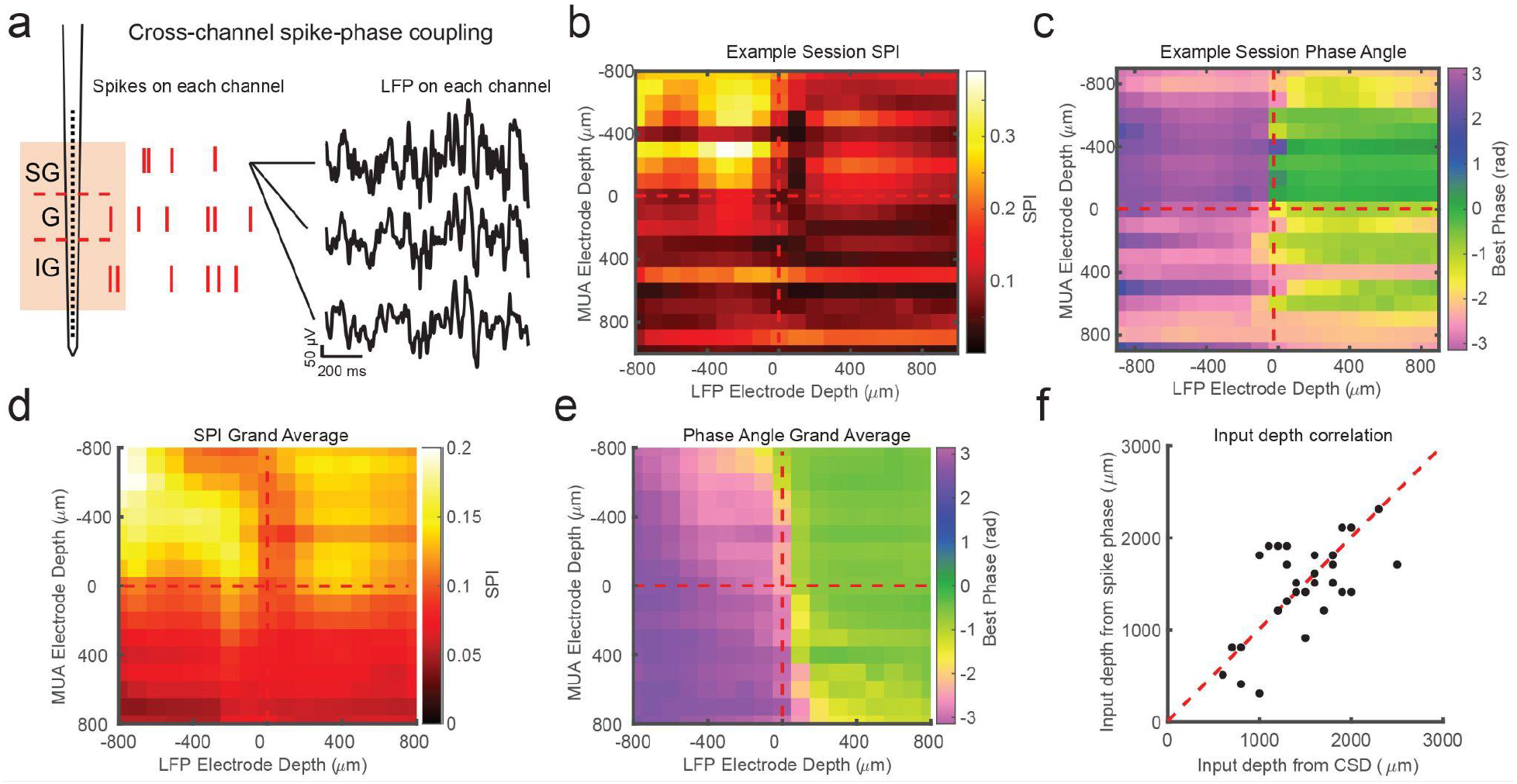
Cross-channel spike-LFP phase coupling reversal correlates with putative input layer. (a) Schematic displaying cross-channel phase coupling analysis. The spike times on each channel are compared against the phase of the LFP on each other channel, yielding a matrix of spike-phase coupling. (b) Example recording session cross-channel SPI values from marmoset MT. Red dashed lines indicate the estimated depth of the bottom of the input layer from CSD analysis. (c) Preferred phase angles for the cross-channel spike-phase relationships in b. Spikes across all channels preferred ±π rad phase angles from LFP measure on superficial and input electrodes, but preferred 0 rad phase angles from LFP measured on deep phase angles. The LFP phase reversal aligns well to the estimate of the input layer from CSD analysis (red dashed line). (d) Grand average cross-channel SPI across all recording sessions in MT, V4, and PFC aligned to the putative input layer from CSD analysis. (e) Grand average of the preferred phase angle for the data in d. (f) Scatter plot shows a significant correlation between the depth of the bottom of the input layer estimated from CSD analysis (x-axis) and the depth of the phase reversal (y-axis; Pearson’s r = 0.64, p = 0.00005).

In order to see if this pattern held across our recordings, we then aligned all of our recordings relative to the boundary that separated input and deep layers (from each recording’s CSD) and computed grand averages for SPI (Figure 3d) and preferred phase angle (Figure 3e). The average pattern across recordings showed the laminar phase reversal was well aligned to the estimate of the input layer from the CSDs across recordings. This was also true when we broke out the recordings to only average across sessions in each cortical area (MT, V4, PFC; Figure S2). To quantify how well the cross-channel LPC and CSD aligned on each individual recording session, we compared the depth of the laminar boundary estimated from CSD analysis to the depth at which the preferred spiking phase angle reversed (Figure 3f). There was a significant correlation between the depth estimated from the CSD and the phase reversal (r = 0.64, p = 0.00005; Pearson’s correlation) and no significant difference between the estimated depth values (mean difference = 271 ± 49 μm S. E. M.; p = 0.98, Wilcoxon signed-rank test).

We next asked whether the observed phase relationship was parameter dependent. We first asked whether or not the specific use of a 5-50 Hz filter or the calculation of GP was necessary to see the laminar phase reversal. To test this we filtered the data in a variety of commonly used frequency bands (4-8 Hz, 8-14 Hz, 15-30 Hz, 30-50 Hz) and calculated phase using the Hilbert Transform (Figure S1). We found that the observed phase reversal was apparent in each case, indicating the results were not dependent on the filter or the method used to compute phase. We also tested whether the results depended on the use of multi-unit spiking activity. We performed the same analysis aligning cross-channel LPC from single-units across recording sessions based on CSD depth (Figure S3). We observed the same relationship between laminar depth and preferred phase angle as when using multi-unit spiking activity. We also compared whether the depth of the phase was dependent on experimental conditions (Figure S4). We separated out different conditions during the same recording session where the electrode placement was unchanged. These included fixation during a visual task, freely viewing natural scene images, freely viewing in total darkness, and fixation during receptive field mapping. The cross-channel LPC revealed qualitatively similar depths for the laminar phase reversal across experimental conditions within the same penetration. Finally, we explored the impact of referencing on the presence of the phase reversal. In our recordings the electrode probe had a reference contact at the base of the shaft. In order to test whether our results depended on the location of the reference contact we re-referenced the LFP data to the shallowest contact, the deepest contact, or a common average reference (Figure S5). The phase reversal persisted under all referencing conditions. These results indicate the phase relationship is a robust feature of columnar recordings and not dependent on a particular set of parameters.

One of the advantages of the LPC method is it can be done without the need for a triggering event, like a stimulus onset. Further, as little as a minute of continuously recorded data is sufficient to recover the phase reversal across channels (Figure S6). We find estimates can be derived from as few as hundreds of multiunit spikes on each channel, although estimates are more reliable when spikes number in the thousands. As a result, depth estimates can be sampled at arbitrary points during recording sessions making this technique sensitive to tracking putative electrode drift over the course of a recording. To demonstrate this, we calculated LPC on sequential 3 minute epochs during the first 15 minutes of an example recording session (Figure S6). We found the depth of the phase reversal moved over the course of 15 minutes. When comparing the depth after the first or last 2 minutes across all recordings, we found a significant difference in the estimated depth of the input layer from the spike-phase reversal (p = 0.005, Wilcoxon signed-rank test) such that the estimate of input layer depth consistently drifted deeper across recording sessions by an average of 132 ± 26 μm (S. E. M.). This change in the depth estimate across the recording is consistent with electrode drift from movement of the probe relative to the cortex as the tissue settles at the start of the recording.

The cross-channel LPC relies on both spikes and LFP, but if a similar phase pattern can be observed in the LFP alone, it may not require spiking activity to detect the laminar boundary. To test this we measured the circular-circular cross correlation between LFP phase on each channel and LFP phase on each other channel. On many recording sessions we found a characteristic phase correlation pattern that seemed to align well with both CSD and LPC depth estimates (Figure 4a-c), and across many recordings the phase correlations within layers could dissociate laminar compartments. However, this was not always the case. Some sessions yielded phase correlation patterns that were noisy and difficult to interpret, although knowing where the boundary was made it easier to identify the characteristic pattern(Figure 4d, e). While, on average, LFP phase correlations can be informative (Figure 4f), the spike-LFP phase relationship appears to provide a more robust measure of contact depth.

**Figure 4.**
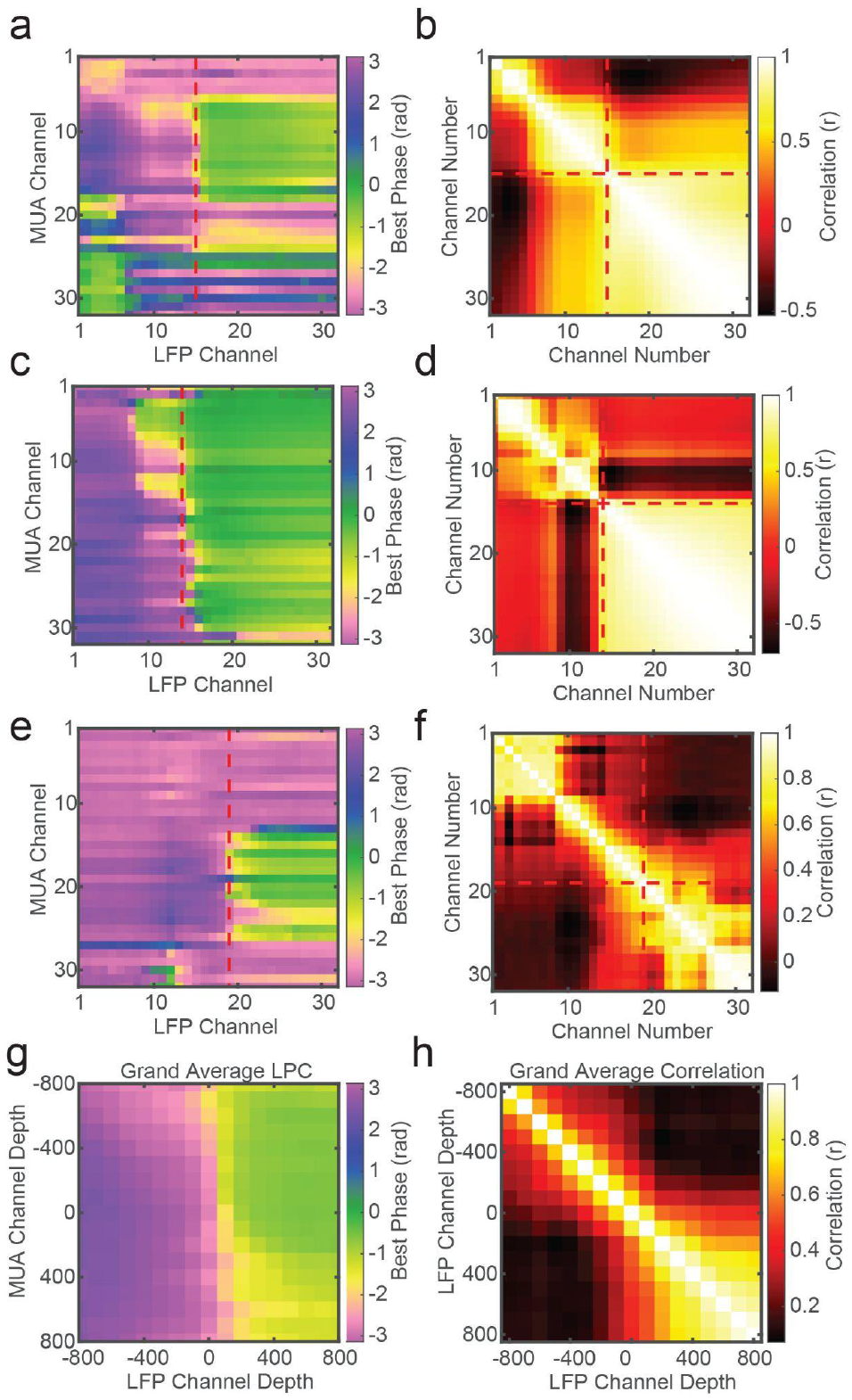
LFP-LFP Phase correlations can also reveal laminar boundaries, but are less consistent. (a) Preferred phase angle across all 32 channels from an example marmoset recording. The red dashed line indicates the CSD estimate of the bottom of the input layer. (b) Circular LFP phase correlation across channels. There is a strong within compartment correlation that aligns with the boundary between channels above and below the input/deep boundary (red dashed lines). (c, d) Same as panels a and b, but for a macaque recording. (e, f) Same as above, but an example of poor phase correlation within compartments despite strong phase reversal alignment with CSD. g, h) Grand averages for preferred phase angle and LFP phase correlations across all recording sessions. Plots were aligned to the putative input/deep layer boundary identified by CSD analysis.

Across our recordings there were some instances (14/34) where there was 200 μm or more disagreement between CSD and LPC estimates for the location of the input layer relative to the probe contacts. Figure 5a shows an example CSD that has the characteristic source-sink pattern used to identify the input layer. This estimate was well aligned to the cross-channel LPC pattern previously described (Figure 5b, c). However, Figure 5d shows an example of a CSD profile with a similar source-sink pattern that is not well aligned to the cross-channel LPC pattern, which shows the characteristic phase reversal 600 μm below the CSD estimate (Figure 5d, e). In order to determine which measurement was more accurate in identifying cortical depth in these cases of disagreement, we sought to use each measure to replicate two findings from the literature that varied with cortical depth.

**Figure 5.**
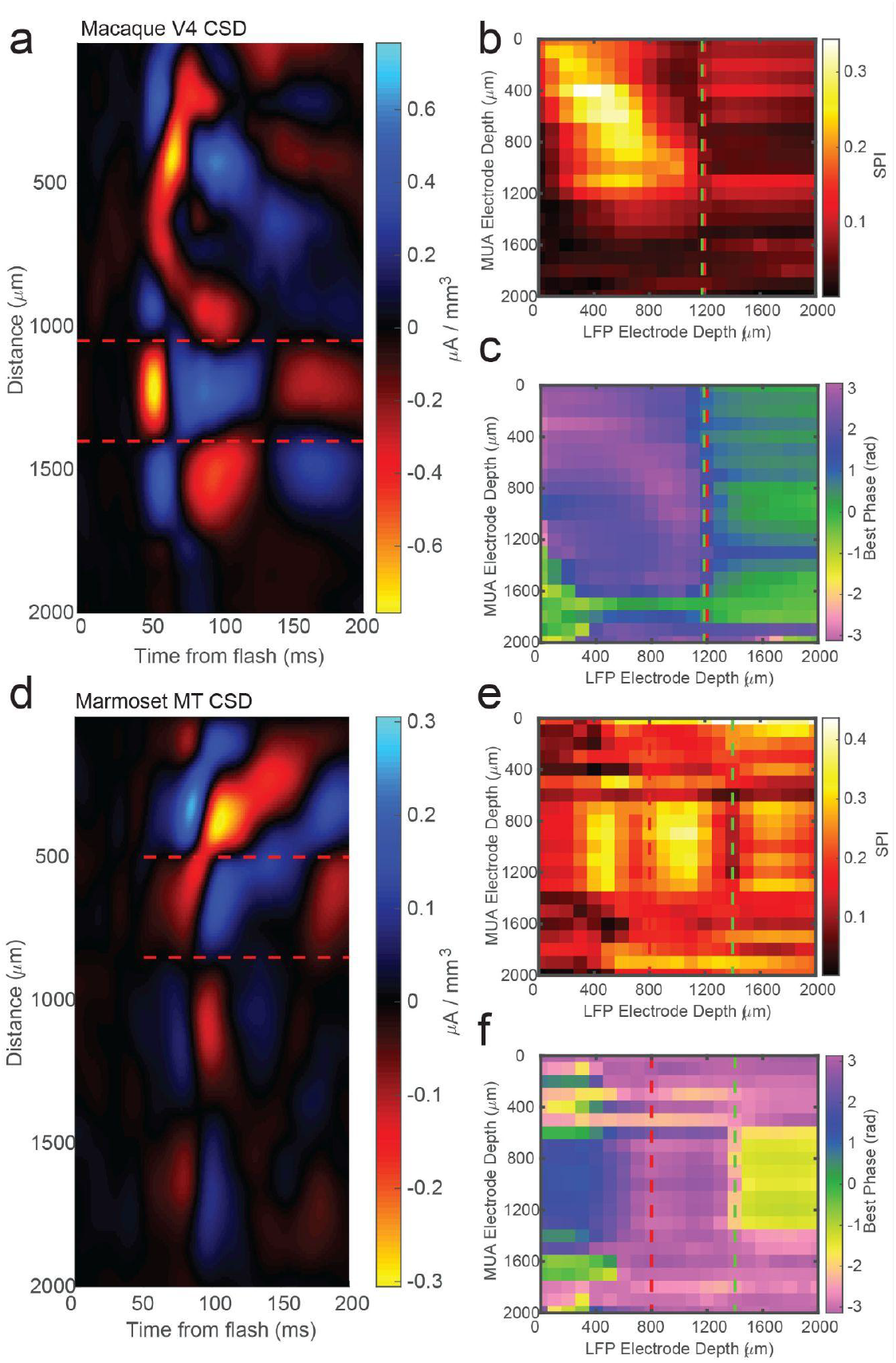
CSD analysis is not always consistent with laminar phase reversal. (a) Example CSD from macaque V4 with estimate of input layer boundaries indicated by red dashed lines. (b) Cross-channel SPI for the same example recording as the CSD in a. Red line is the CSD estimate of the bottom of the input layer. Green line is the estimate from the phase reversal. (c) Cross-channel spike-phase pattern for same recording as in a and b. The phase reversal is well aligned to the CSD estimate of the input layer. (d) CSD from a different example recording session in marmoset MT. Laminar depth estimated the same as in a. (e, f) Cross-channel SPI and phase angle as in b and c. The CSD and phase reversal estimates do not align.

First, previous work in multiple cortical areas has shown that the superficial layers have lower firing rates as compared to the input and deep layers (Lakatos et al., 2005; de Kock and Sakmann, 2009; Haegens et al., 2015; Leszczyński et al., 2020). We identified recording sessions where the CSD and LPC estimate of input depth differed by more than 200 μm (N = 14 sessions) and we measured the average firing rate as a function of depth relative to either the CSD or LPC estimate of the input layer (Figure 6a). We found that on these sessions, the CSD aligned firing rates failed to recapitulate the finding of higher firing rates in the input and deeper layers (p = 0.09, WIlcoxon signed-rank test). Conversely, we did find significantly stronger input firing rates when aligned to the LPC estimate (p = 0.0003, Wilcoxon signed-rank test), suggesting the LPC estimate on these sessions was better aligned than the CSD to the ground truth.

**Figure 6.**
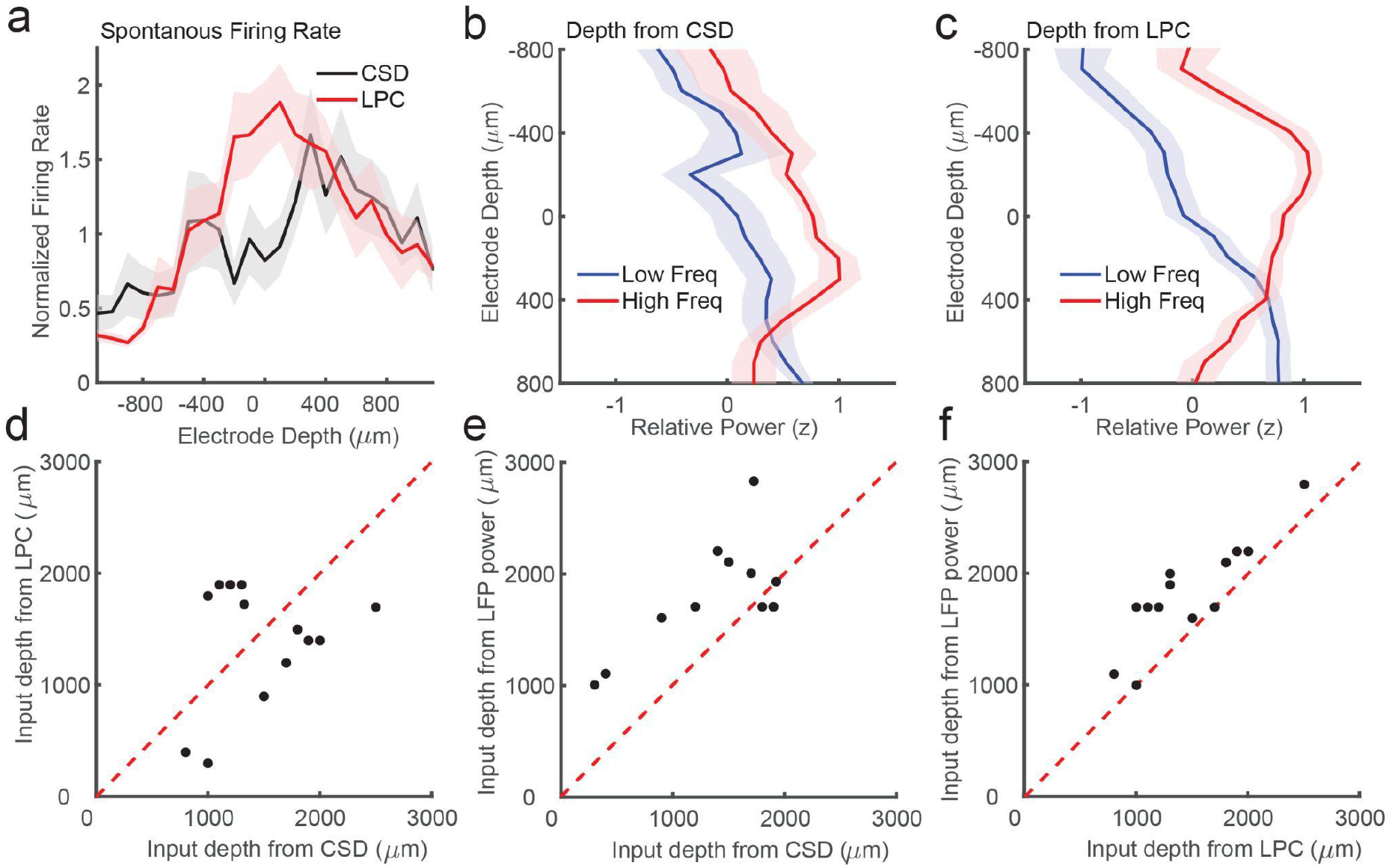
Laminar phase coupling pattern better replicates spike rate and LFP power dissociations than CSD when the two measures disagree. (a) Normalized spontaneous firing rates across sessions when CSD and LPC disagreed as a function of contact depth by more than 200 μm (N = 14). LPC captures the expected higher spontaneous firing rates around the input layer. (b) Mean z-scored LFP power across sessions in lower frequencies (8-30 Hz, blue line) and higher frequencies (65-100 Hz, red line) as a function of electrode depth referenced to the CSD estimate for sessions. (c) Same as in a, but referenced to the depth estimate from the LPC phase reversal across sessions. The laminar power relationship is more pronounced when using the phase reversal instead of CSD estimate. (d) Scatter plot showing the disagreement in the depth of the input layer estimated from CSD (x-axis) and the depth from LPC (y-axis) in this subset of recording sessions. (e) Scatter plot showing the alignment of the CSD input depth (x-axis) cross-over in LFP power (y-axis). (f) Same as e, but for LPC. The LPC depth measure was more correlated with the LFP power reversal than the CSD measure (Pearson’s r = 0.65 v 0.87 CSD v LPC).

We next examined a second previously reported finding that varied with laminar depth. Previous groups have reported an inversion in the spectral content of the LFP across layers, with the superficial layers exhibiting more high frequency power (e.g. 80-200 Hz) and the deep layers exhibiting more low frequency power (8-20 Hz)(Maier et al., 2010, 2011). The crossover between the power in higher and lower frequencies was reported to occur around the input layers defined from CSD measurements(Bastos et al., 2018). To test whether we could identify a similar relationship in these ambiguous recordings, and whether that relationship was stronger when aligned to CSD or LPC estimates of depth, we calculated low and high frequency power on each channel in each session and aligned them to each depth estimate (Figure 6b, c). The depth estimate from the LPC measure was significantly correlated with the high/low power inversion (Pearson’s r = 0.87, p = 0.00005) whereas the CSD estimate was more weakly, although still significantly correlated (Pearson’s r = 0.65, p = 0.012), indicating that on the recording sessions where there was some disparity between the CSD and LPC estimate (Figure 6d), the LPC better captured the power inversion than the CSD measure (Figure 6e, f). These results suggest that when noise or error in assigning laminar compartments based on CSD analysis occur, the LPC phase reversal is a more reliable measure of laminar depth.

## Discussion

The cortical column is one of the fundamental computational circuits in the brain. In order to understand the function of disparate cortical areas and the contribution various cell-types play in processing information through the columnar circuit, it is often necessary to identify and segregate the responses of neuronal populations based on their position in the layers of the cortex. Traditionally, for electrophysiological measurements, this has been achieved using CSD analysis of sensory-evoked responses. However, CSD analysis requires averaging across repeated discrete sensory events that may be difficult to generate in less well studied cortical areas where the response selectivity is not apparent. While CSDs can also be calculated from intrinsic events such as up/down state transitions (Senzai et al., 2019) or bursts of oscillatory power (Bollimunta et al., 2008), it is less clear how reliable their patterns reflect the laminar organization of the underlying circuitry with respect to distinct anatomical boundaries—although methods have been proposed to recover spatial information from spontaneous activity(Chand and Dhamala, 2014). Further, because CSDs are measured across electrodes, a single noisy channel corrupts the contribution of the channel above and below potentially leading to an errant assignment of cortical layer boundaries. As histological approaches to recover individual recording tracts can be impractical, particularly in experiments in non-human primate where multiple recordings in the same area are the norm, it can be difficult to validate CSD measures or recover laminar information.

Here we describe an alternative estimate of laminar boundary, laminar-phase coupling (LPC), that can supplement or in some cases replace CSD analysis. Using laminar boundaries defined by CSD, we find that spike-phase coupling inverts at the boundary between the input and deep layers of the cortex. This reversal provides a reliable and robust measure of the depth of linear array electrodes in cortex across a variety of analysis parameters and experimental conditions. This measure is robust to different LFP filter settings with either single- and multi-unit data and can be applied to any arbitrary recording epoch so long as there are sufficient spikes across electrodes. Failing that, patterns of phase-phase correlation across channels may also help identify laminar boundaries. The ability to identify cortical depth across linear electrodes quickly and robustly permits the online identification of electrode positions relative to cortical layers and the tracking of electrode drift as the cortex settles following electrode penetration.

While the number of spikes necessary to recover the spike-phase inversion can be counted in the hundreds, the LPC measurement is more reliable in epochs in which thousands of spikes have occurred. It would be convenient if LFP phase alone were sufficient to identify cortical layers as this would eliminate the requirement for measuring multi-unit spiking on multiple electrodes, and indeed we find there are occasions where correlations in LFP phase are sufficient to provide strong evidence of the location of laminar boundaries. This is consistent with previous reports of dissociable patterns of LFP coherence between superficial and deep domains that show particular separation at the bottom of the input layer(Maier et al., 2010). However, we find the reliability of this pattern of LFP phase correlation varies across the recordings in our dataset, although the reason why is unclear. We do find measures of cortical depth are improved when LFP phase is combined with the preferred timing of spiking activity across the cortical layers. However, if multi-unit data is only weakly apparent in a linear array recording, depth information might still be recovered from LFP phase relationships across channels without requiring CSD analysis.

One interesting observation is that the superficial and input layers of cortex show stronger spike-LFP coupling than the deeper layers. Why might this be the case? It may be that the LFP is largely reflecting the shared synaptic inputs in the numerous connections in these cortical layers(Buzsáki et al., 2012). This is consistent with measurements of correlated variability in V2 where spike-count correlations were observed to be stronger in the superficial and input layers and weaker in the deep layers(Smith et al., 2013), or in V4 where the strongest correlations were observed in the input layers relative to the superficial or deep layers(Nandy et al., 2017). A different pattern has been observed in V1, where the superficial and deep layers showed stronger spike-count correlations and in the input layers showed weaker correlations(Hansen et al., 2012). One might predict that a different pattern of spike-phase coupling would occur in V1, although the contribution of an anesthetized state is unclear.

We find reliable patterns across V4, MT, and PFC, and across two species of monkey performing visually guided tasks. Our findings may be limited to these conditions, as we do not yet know if the same observations hold in cortical areas with non-sensory responses, under conditions of sleep or anesthesia, or in other species. The answer may rely on what generates the observed pattern of spike-LFP phase relationship. While our purpose here is to provide an empirically determined alternative or supplement to CSD analysis, we can speculate about why this observed LPC reversal occurs at the transition from input to deep layers. One possibility stems from the hypothesis that the predominantly radial architecture of cortical fibers between superficial and deep layers forms an electrical dipole that spans across the layers of the cortex(Buzsáki et al., 2012). This hypothesis underlies the generation of electrical fields parallel to the dipole measured with EEG recordings at the scalp, and the generation of magnetic fields orthogonal to the dipole measured in MEG recordings. The prediction, therefore, is that the observed spike-phase reversal occurs due to the sources of LFP phase inverting across a dipole generated by the structure of the parallel radial processes of pyramidal cells that arborize in the superficial layers (however see (Riera et al., 2012)). Future experiments dissociating the laminar contribution of cell populations to the LFP at different cortical depths may reveal the degree to which the observed phase reversal is due to the dipole hypothesis or some other circuit mechanism. While the network mechanism responsible for the observed phase reversal remains unclear, our results indicate the reversal is well aligned to the boundary that separates the input and deep layers in multiple cortical areas.

The LPC method for estimating laminar depth described here could be a useful supplement to CSD analysis as it provides a source of cross-validation for potentially noisy or ambiguous CSD patterns. It could also serve as an alternative source of laminar information when CSD analysis is impractical given experimental limitations. The advantages of the LPC method are that it is fast and simple, making it well suited to online depth estimation from unsorted multiunit spiking activity. LPC can be estimated from ongoing activity in a variety of experimental conditions, cortical areas, and analysis parameters, and may help relate the function of neural populations to the fundamental computation of information processing in the columnar cortical circuit.

---

**Figure S1.**
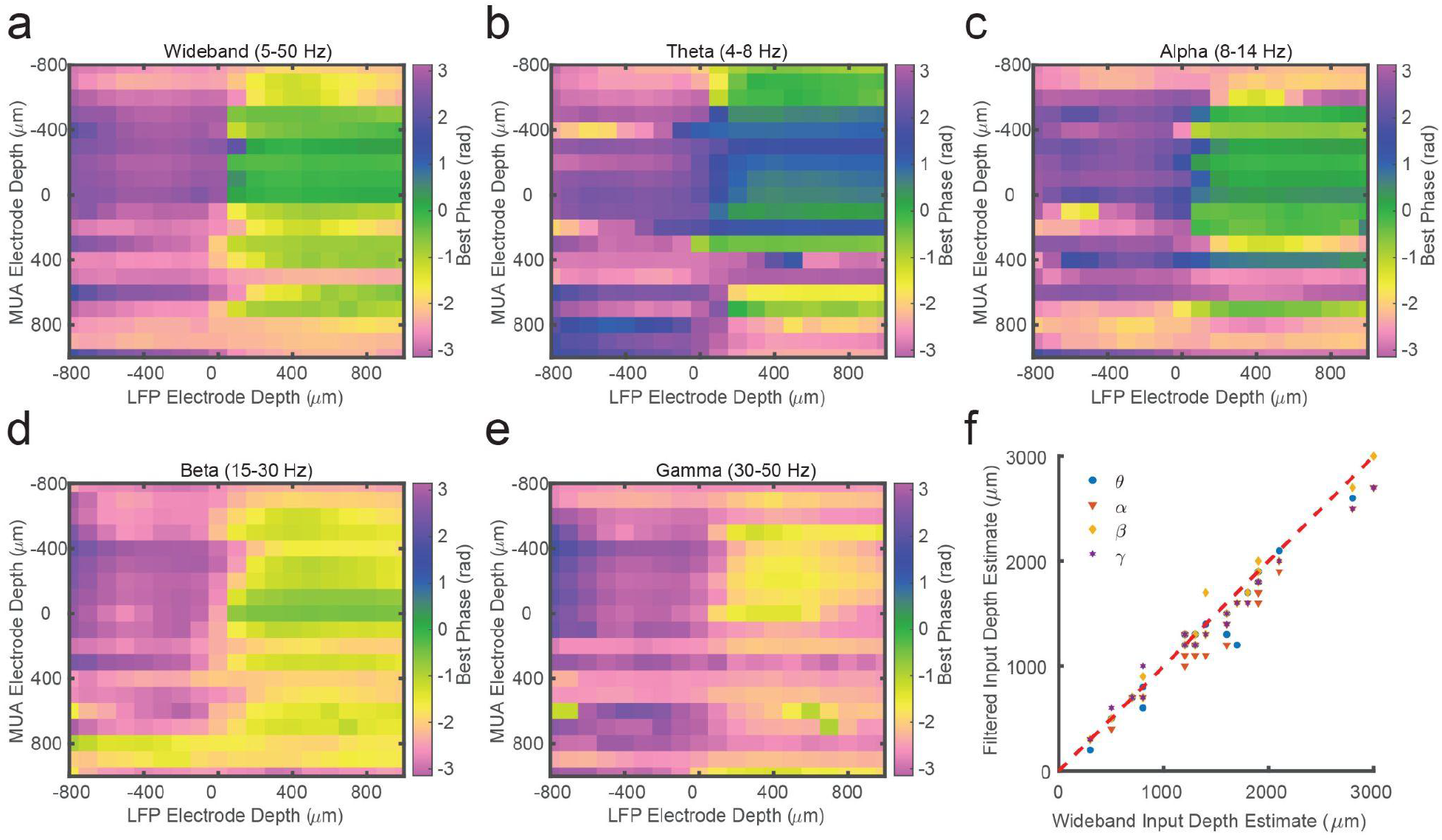
Laminar spike-phase pattern does not depend on filtering. (a) Example cross-channel LPC from marmoset MT after filtering LFP from 5-50 Hz calculated using GP. (b-e) Same as in a, but after filtering in theta (4-8 Hz), alpha (8-14 Hz), beta (15-30 Hz), or low gamma (30-50 Hz) frequency bands and measuring phase with the Hilbert transform. The LPC reversal occurs at the same depth regardless of filter bandwidth. (f) Scatter plot showing the lack of dispersion from unity between depth estimates made using the 5-50 Hz filter or each of the tested narrowband filters.

**Figure S2.**
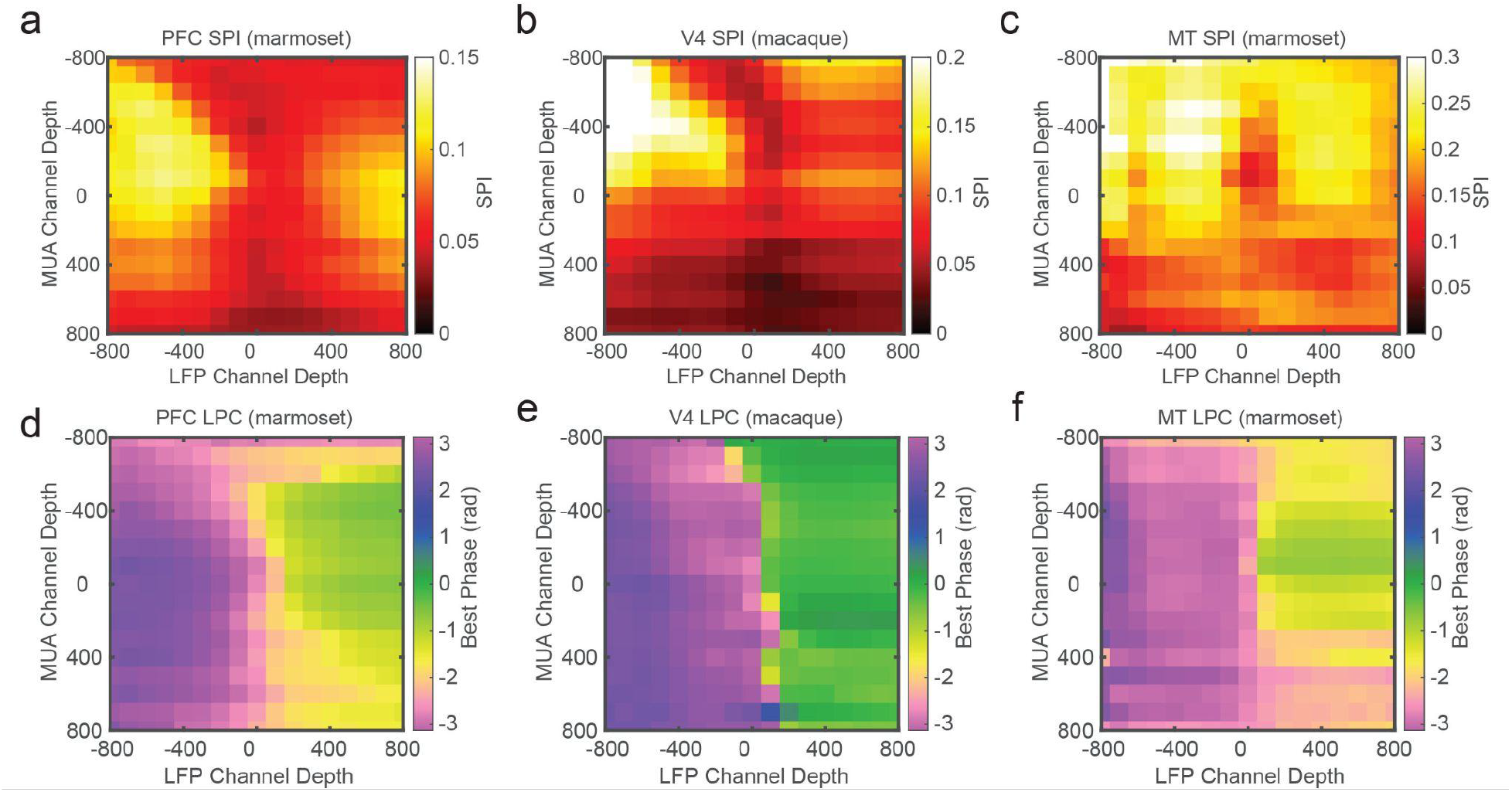
Laminar spike-phase reversal consistent across cortical areas. (a-c) Grand average cross-channel SPI measurements for sessions recorded in marmoset PFC (a), macaque V4 (b), and marmoset MT (c). The depth measurements are relative to the estimate of the bottom of the input layer from CSD analysis. (d-f). Grand average cross-channel preferred spike-phase angles for the cortical areas as in a-c. The phase reversal is well aligned to the putative input layer estimated from CSD analysis.

**Figure S3.**
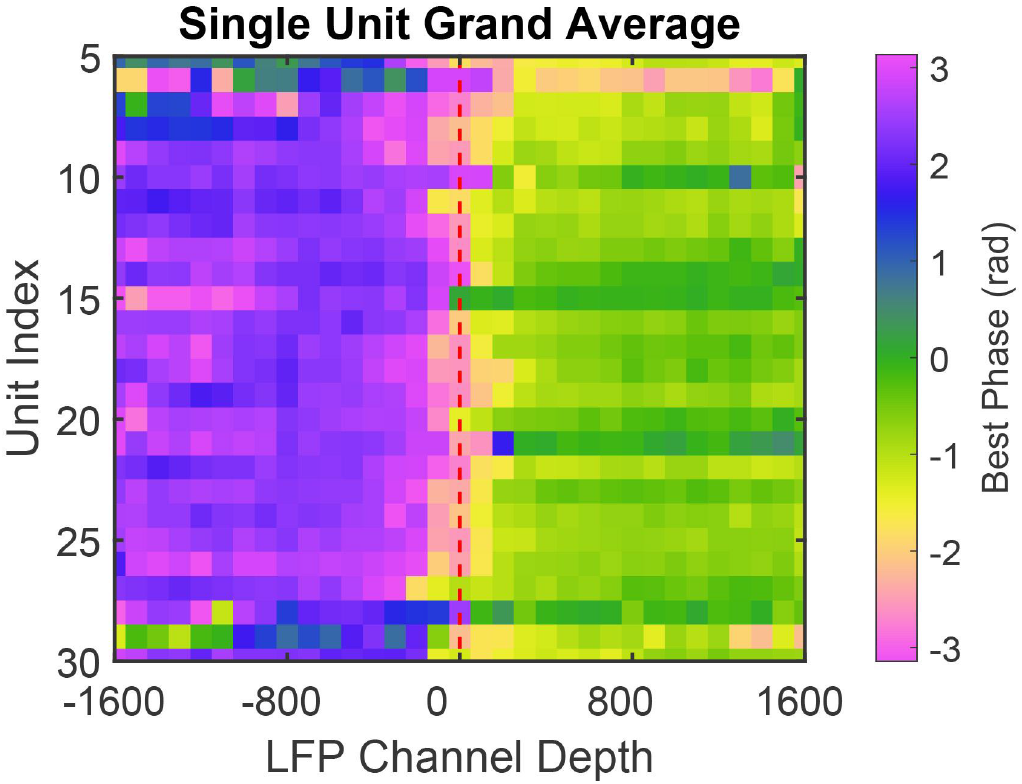
Phase reversal is apparent when using well isolated single units instead of MUA. LPC grand average across all sessions for well isolated single units. Units were aligned across sessions relative to their index from the median index. The use of single-unit spiking instead of MUA produces qualitatively similar results.

**Figure S4.**
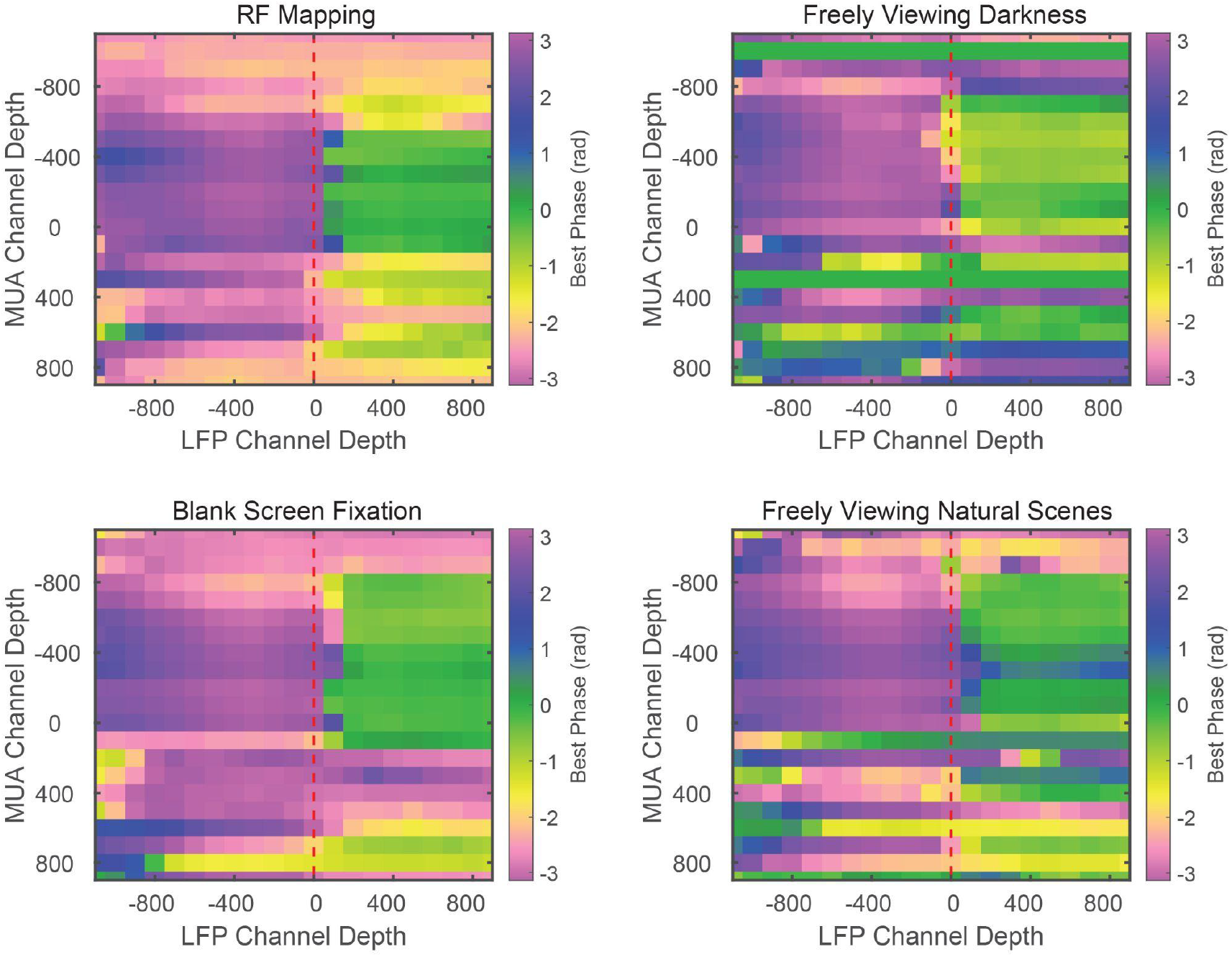
Phase relationship does not depend on task conditions. Example cross-channel LPC measurements for different experimental conditions during the same recording session. The spike-phase reversal is consistent across receptive field mapping, free viewing in total darkness, fixation on a blank screen during a visual detection task, or freely viewing naturalistic images. Example data from marmoset MT.

**Figure S5.**
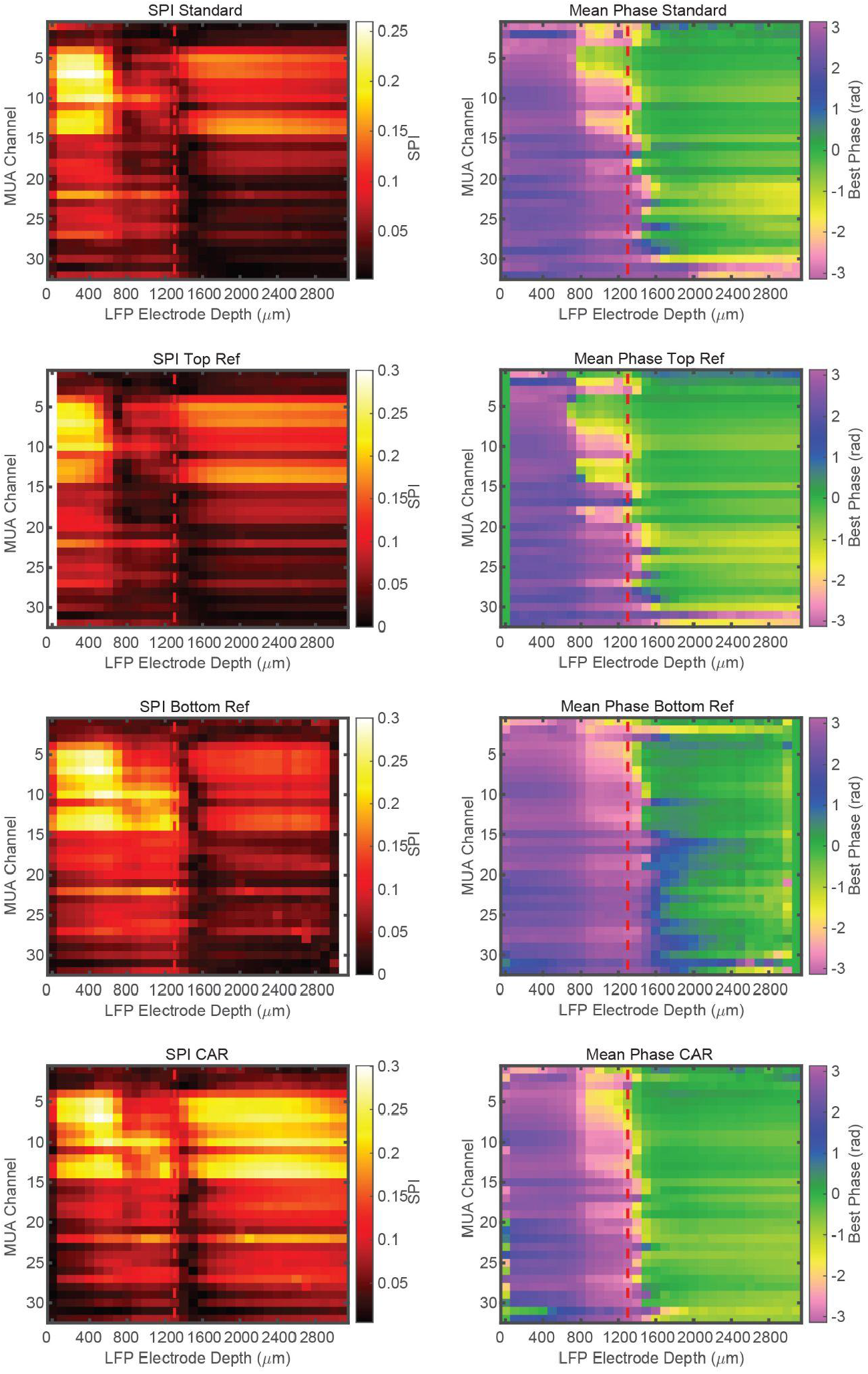
Phase relationship is not dependent on referencing. Cross-channel LPC measurements from the same recording session with the integrated reference contact, re-referencing to the top most contact, the bottom most contact, or a common average reference (CAR) from top to bottom respectively. The presence of the laminar phase reversal is not dependent on referencing. Example data from macaque V4.

**Figure S6.**
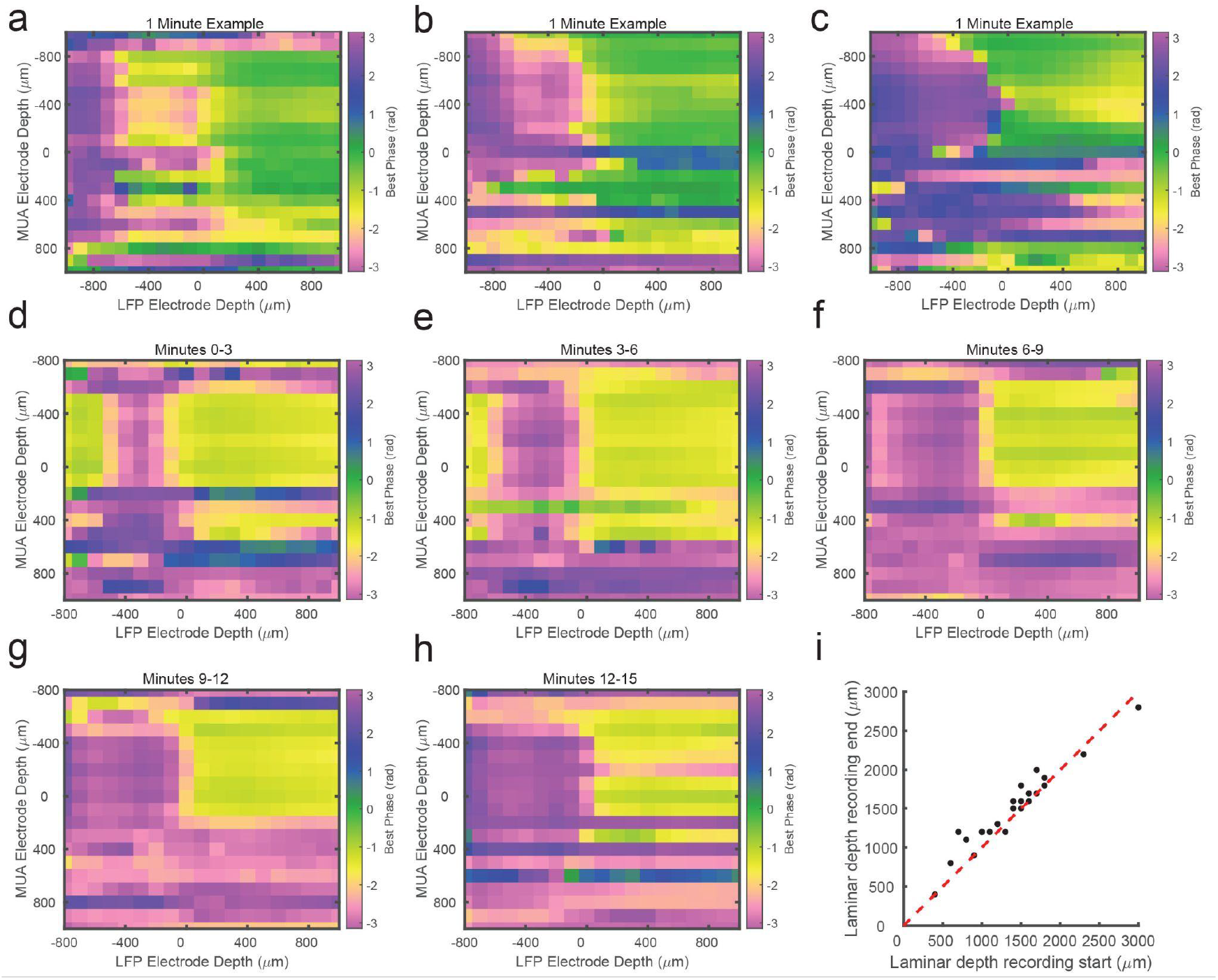
Cross-channel LPC can be estimated over short periods. (a-c) Cross-channel LPC measurements from 3 example sessions generated from 1 minute of recorded data. The phase reversal in these sessions is well aligned to the estimate from the entire recording. (d) LPC measurement from the first 3 minutes of a different linear probe recording. (e-h) LPC measurements of the subsequent 3 minutes of the same recording in d. The location of the phase reversal drifts over the course of the first 15 minutes of the recording. (f) Scatter plot comparing the estimated depth of the phase reversal based on the first (x-axis) or last (y-axis) 2 minutes of each recording session. Drift of 100-300 μm is consistently observed across recordings.

## Materials and Methods

### Surgical Approach

Data from two male marmoset monkeys *(Callithrix jacchus)* and two male rhesus macaques (*Macaca mulatta*) were used in this study. Data from the macaques were previously published in (Franken and Reynolds, 2021). Macaque surgical procedures have been described before in (Nandy et al., 2017). Similar techniques were used in the marmoset monkeys. Monkeys were surgically implanted with headposts for head stabilization and eye tracking using cranial screws and dental acrylic or cement. In subsequent surgical sessions a titanium recording chamber was installed in a craniotomy made over area MT or PFC in one marmoset respectively, or area V4 in both macaques, according to stereotactic coordinates. The dura mater within the chamber was removed, and replaced with a silicone-based optically clear artificial dura, establishing an optical window over the cortex. All surgical procedures were performed with the monkeys under general anesthesia in an aseptic environment in accordance with the recommendations in the Guide for the Care and Use of Laboratory Animals of the National Institutes of Health. All experimental methods were approved by the Institutional Animal Care and Use Committee (IACUC) of the Salk Institute for Biological Studies and conformed with NIH guidelines.

### Electrophysiological Recordings

Electrode voltages were recorded from a 32 channel linear silicone electrode array (Atlas Neuroengineering; Leuven, Belgium) connected to an Intan RHD2000 USB interface board (Intan Technologies LLC, Los Angeles, USA) controlled by a Windows computer. The probe was inserted through the artificial dura using a hydraulic microdrive mounted on the chamber using an adjustable *x*–*y* stage (MO-972A, Narashige, Japan). The probe was lowered until spiking and LFP could be observed on most of the electrode contacts. Then, the probe was retracted typically by several 100 μm to ease dimpling of the cortex. Data were sampled at 30 kHz from all channels. Neural data was broken into two streams for offline processing of spikes (single-unit and multi-unit activity) and LFPs. Spike data was high-pass filtered at 500 Hz and candidate spike waveforms were defined as exceeding 4 standard deviations of a sliding 1 second window of ongoing voltage fluctuations. Sorted units were classified as single- or multi-units and single units were validated by the presence of a clear refractory period in the autocorrelogram. LFP data was low-pass filtered at 300 Hz and down-sampled to 1000 Hz.

### Behavioral tasks

Marmosets were trained to enter a custom-built marmoset chair that was placed inside a faraday box with an LCD monitor (ASUS VG248QE) at a distance of 40 cm. The monitor was set to a refresh rate of 100 Hz and gamma corrected with a mean gray luminance of 75 candelas/m^2^. For the macaques visual stimuli were presented using a LED projector, back-projected on a rear-projection screen that was positioned at a distance of 52 cm from the animal’s eyes (PROPixx, VPixx Technologies, Saint-Bruno, Canada). The marmosets and macaques were headfixed by a headpost for all recordings. Eye position was measured with an IScan CCD infrared camera. The MonkeyLogic software package developed in MATLAB (https://www.brown.edu/Research/monkeylogic/; https://monkeylogic.nimh.nih.gov/)(Asaad et al., 2013) was used for stimulus presentation, behavioral control, and recording of eye position. Digital and analog signals were coordinated through National Instrument DAQ cards (NI PCI6621) and BNC breakout boxes (NI BNC2090A).

Ongoing data while marmosets and macaques performed a variety of tasks and viewing conditions were used in this study. These conditions included fixation of a fixation point, receptive field mapping, freely viewing natural images, freely viewing in a darkened room, and performing previously described visual tasks (marmoset: (Davis et al., 2020), macaque: (Franken and Reynolds, 2021)). Not all tasks were included in each monkey’s experimental battery. All data were collapsed across applicable conditions as the results did not depend on sensory conditions (Figure S4). 6 recording sessions from V4 in each macaque, 8 recording sessions in marmoset MT, and 14 recording sessions from marmoset PFC were used. Sessions were only included if CSD analysis revealed a recognizable source-sink pattern consistent.

### Current-Source Density Analysis

A CSD mapping procedure on evoked local field potentials (LFP) was used to estimate the laminar position of recorded channels(Nandy et al., 2017; Franken and Reynolds, 2021). Mapping stimuli varied across recording locations. For macaque recordings in V4, monkeys maintained fixation while dark gray ring stimuli were flashed (32 ms stimulus duration, 94% luminance contrast, sized and positioned to fall within the cRF of the probe position). For marmoset recordings in MT, monkeys maintained fixation while drifting Gabor stimuli (spatial frequency = 0.5 cycles per degree; temporal frequency = 10 cycles per second, 50% luminance contrast) were presented. For marmoset recordings in PFC, monkeys maintained fixation while full field 100% luminance flashes (background luminance 0.5 candelas/m^2^; 20 ms flash at 150 candelas/m^2^) were presented. The CSD was calculated as the second spatial derivative of the stimulus-triggered LFP and visualized as spatial maps after smoothing using bicubic (2D) interpolation (MATLAB function *interp2* with option *cubic*), although the laminar analysis did not critically depend on this particular method of smoothing. Red regions depict current sinks, blue regions depict current sources. We identified the earliest current sink as the input layer. By comparing this position with the range of contacts in the input layer, we could locate channels to superficial, input, or deep layers.

### Generalized Phase

We calculated Generalized Phase (GP) as described previously(Davis et al., 2020). The purpose of GP is to mitigate the breakdown of the analytic signal representation for spectrally broad signals. As an initial step in the GP representation, then, we filter the signal within a wide bandpass (i. e. 5-50 Hz; 4^th^-order zero-phase Butterworth filter), excluding low-frequency content that contributes to origin offsets in the complex plane that distort the estimate of phase angles for higher frequency signals. We then use the single-sided Fourier transform approach(Johansson, 1999; Marple, 1999) on the wideband signal and compute phase derivatives as finite differences, which are calculated by multiplications in the complex plane(Feldman, 2011/4; Muller et al., 2014, 2016). High-frequency intrusions appear in the analytic signal representation as complex riding cycles(Feldman, 2011/4), which manifest as periods of negative frequencies in the analytic signal representation. As a secondary step we then numerically detect these complex riding cycles (*N_c_* points of negative frequency) and utilize shape-preserving piecewise cubic interpolation on the next *2N_c_* points following the detected negative frequency epoch. The resulting representation captures the phase of the largest fluctuation on the recording electrode at any moment in time, without the distortions due to the large, low-frequency intrusions or the smaller, high-frequency intrusions characteristic of the *1/f*-type fluctuations in cortical LFP(Pereda et al., 1998; Linkenkaer-Hansen et al., 2001; Milstein et al., 2009).

### Laminar analyses

The degree of spike-phase coupling was measured as the mean resultant vector length for the LFP phase distribution collected at the time of observed spikes. This measure was calculated using the circ_r function in the Circular Statistics Toolbox for MATLAB(Berens, 2009). The mean phase angle of the spike-phase distribution was calculated using the circ_mean function in the Circular Statistics Toolbox. To generate the cross-channel LPC, the phase of the LFP on each channel was collected for the spike train on each channel. The mean spike-phase angle for each combination of spike and LFP channel was plotted, and the channel that best separated the preferred phase angle was selected by eye as the boundary between the input and deep layers. This process was done blind to the CSD estimate of laminar depth.

For laminar analyses comparing firing rates and LFP power across cortical layers N = 14 sessions were selected based on a difference in laminar alignment between CSD and LPC analyses of more than 200 μm. Each channel was aligned to the estimate of the boundary between the input and deep layers in each session based on either the bottom of the earliest current sink in the CSD or the channel preceding the change in the pattern of LPC phase coupling. Mean spike rates at each channel depth were normalized to the mean spike rate on all channels in each session. LFP power in low (8-30 Hz) and high (65-100 Hz) frequency bands were calculated by taking the average power spectral density in the frequency range of interest at each electrode depth normalized by the average power spectral density in both frequency ranges across all channels. The values were then z-scored across channels.

Statistical differences were determined by Wilcoxon signed-rank tests for pairwise differences in the distributions across superficial, input, or deep layers within CSD or LPC conditions, as well as across CSD and LPC conditions. Differences in circular distributions were computed by Watson-Williams test of homogeneity of means. Circular correlations were calculated using the circ_corrcc function for circular-circular correlations and the circ_corrcl function for circular-linear correlations in the MATLAB Circular Statistics Toolbox.

## Author Contributions

Conceptualization, Z.W.D., N.M.D., T.F.; Data Curation, Z.W.D, N.M.D, T.F.; Formal Analysis, Z.W.D; Funding Acquisition, Z.W.D., L.M., J.R.; Investigation, Z.W.D.; Methodology, Z.W.D., N.F.D., T.F., L.M.; Supervision, L.M., J.R.; Visualization, Z.W.D.; Writing - original draft, Z.W.D., N.M.D., T.F., L.M., J.R.; Writing - review and editing, Z.W.D., N.M.D., T.F., L.M., J.R.

## Data Availability

The data that support the findings of this study are available from the corresponding author upon reasonable request.

## Code Availability

An open-source code repository for measuring LPC is available on http://github.com/zwdsalk/LaminarPhaseCoupling

## Acknowledgments

Funding: Gatsby Charitable Foundation, the Fiona and Sanjay Jha Chair in Neuroscience, the Canadian Institute for Health Research, the Swartz Foundation, NIH grants R01-EY028723, T32 EY020503-06, P30 EY019005, K99/R00 EY031795 and BrainsCAN at Western University through the Canada First Research Excellence Fund (CFREF).

## References

Asaad WF, Santhanam N, McClellan S, Freedman DJ (2013) High-performance execution of psychophysical tasks with complex visual stimuli in MATLAB. J Neurophysiol 109:249–260.

Bastos AM, Loonis R, Kornblith S, Lundqvist M, Miller EK (2018) Laminar recordings in frontal cortex suggest distinct layers for maintenance and control of working memory. Proc Natl Acad Sci U S A 115:1117–1122.

Berens P (2009) CircStat: AMATLABToolbox for Circular Statistics. Journal of Statistical Software 31 Available at: http://dx.doi.org/10.18637/jss.v031.i10.

Bode-Greuel KM, Singer W, Aldenhoff JB (1987) A current source density analysis of field potentials evoked in slices of visual cortex. Exp Brain Res 69:213–219.

Bollimunta A, Chen Y, Schroeder CE, Ding M (2008) Neuronal mechanisms of cortical alpha oscillations in awake-behaving macaques. J Neurosci 28:9976–9988.

Buzsáki G, Anastassiou CA, Koch C (2012) The origin of extracellular fields and currents--EEG, ECoG, LFP and spikes. Nat Rev Neurosci 13:407–420.

Chand GB, Dhamala M (2014) Spectral factorization-based current source density analysis of ongoing neural oscillations. J Neurosci Methods 224:58–65.

Davis ZW, Muller L, Martinez-Trujillo J, Sejnowski T, Reynolds JH (2020) Spontaneous travelling cortical waves gate perception in behaving primates. Nature 587:432–436.

Davis ZW, Muller L, Reynolds JH (2022) Spontaneous Spiking Is Governed by Broadband Fluctuations. J Neurosci 42:5159–5172.

de Kock CPJ, Sakmann B (2009) Spiking in primary somatosensory cortex during natural whisking in awake head-restrained rats is cell-type specific. Proc Natl Acad Sci U S A 106:16446–16450.

Destexhe A, Contreras D, Steriade M (1999) Spatiotemporal analysis of local field potentials and unit discharges in cat cerebral cortex during natural wake and sleep states. J Neurosci 19:4595–4608.

Dotson NM, Salazar RF, Gray CM (2014) Frontoparietal correlation dynamics reveal interplay between integration and segregation during visual working memory. J Neurosci 34:13600–13613.

Eeckman FH, Freeman WJ (1990) Correlations between unit firing and EEG in the rat olfactory system. Brain Res 528:238–244.

Einevoll GT, Kayser C, Logothetis NK, Panzeri S (2013) Modelling and analysis of local field potentials for studying the function of cortical circuits. Nat Rev Neurosci 14:770–785.

Esghaei M, Daliri MR, Treue S (2018) Attention decouples action potentials from the phase of local field potentials in macaque visual cortical area MT. BMC Biol 16:86.

Feldman M (2011/4) Hilbert transform in vibration analysis. Mech Syst Signal Process 25:735–802.

Franken TP, Reynolds JH (2021) Columnar processing of border ownership in primate visual cortex. Elife 10 Available at: http://dx.doi.org/10.7554/eLife.72573.

Godlove DC, Maier A, Woodman GF, Schall JD (2014) Microcircuitry of agranular frontal cortex: testing the generality of the canonical cortical microcircuit. J Neurosci 34:5355–5369.

Gratiy SL, Devor A, Einevoll GT, Dale AM (2011) On the estimation of population-specific synaptic currents from laminar multielectrode recordings. Front Neuroinform 5:32.

Haegens S, Barczak A, Musacchia G, Lipton ML, Mehta AD, Lakatos P, Schroeder CE (2015) Laminar Profile and Physiology of the α Rhythm in Primary Visual, Auditory, and Somatosensory Regions of Neocortex. J Neurosci 35:14341–14352.

Haegens S, Nácher V, Luna R, Romo R, Jensen O (2011) α-Oscillations in the monkey sensorimotor network influence discrimination performance by rhythmical inhibition of neuronal spiking. Proc Natl Acad Sci U S A 108:19377–19382.

Hansen BJ, Chelaru MI, Dragoi V (2012) Correlated variability in laminar cortical circuits. Neuron 76:590–602.

Happel MFK, Jeschke M, Ohl FW (2010) Spectral integration in primary auditory cortex attributable to temporally precise convergence of thalamocortical and intracortical input. J Neurosci 30:11114–11127.

Johansson M (1999) The hilbert transform. Mathematics Master’s Thesis Växjö University, Suecia Disponible en internet: http://w3msivxuse/exarb/mj_expdf, consultado el 19 Available at: http://www.fuchs-braun.com/media/d9140c7b3d5004fbffff8007fffffff0.pdf

Lakatos P, Chen C-M, O’Connell MN, Mills A, Schroeder CE (2007) Neuronal oscillations and multisensory interaction in primary auditory cortex. Neuron 53:279–292.

Lakatos P, Shah AS, Knuth KH, Ulbert I, Karmos G, Schroeder CE (2005) An oscillatory hierarchy controlling neuronal excitability and stimulus processing in the auditory cortex. J Neurophysiol 94:1904–1911.

Leszczyński M, Barczak A, Kajikawa Y, Ulbert I, Falchier AY, Tal I, Haegens S, Melloni L, Knight RT, Schroeder CE (2020) Dissociation of broadband high-frequency activity and neuronal firing in the neocortex. Sci Adv 6:eabb0977.

Linkenkaer-Hansen K, Nikouline VV, Palva JM, Ilmoniemi RJ (2001) Long-range temporal correlations and scaling behavior in human brain oscillations. J Neurosci 21:1370–1377.

Maier A, Adams GK, Aura C, Leopold DA (2010) Distinct superficial and deep laminar domains of activity in the visual cortex during rest and stimulation. Front Syst Neurosci 4 Available at: http://dx.doi.org/10.3389/fnsys.2010.00031.

Maier A, Aura CJ, Leopold DA (2011) Infragranular sources of sustained local field potential responses in macaque primary visual cortex. J Neurosci 31:1971–1980.

Marple L (1999) Computing the discrete-time“analytic” signal via FFT. IEEE Trans Signal Process 47:2600–2603.

Milstein J, Mormann F, Fried I, Koch C (2009) Neuronal shot noise and Brownian 1/f2 behavior in the local field potential de la Prida LM, ed. PLoS One 4:e4338.

Mitzdorf U (1985) Current source-density method and application in cat cerebral cortex: investigation of evoked potentials and EEG phenomena. Physiol Rev 65:37–100.

Mitzdorf U (1987) Properties of the evoked potential generators: current source-density analysis of visually evoked potentials in the cat cortex. Int J Neurosci 33:33–59.

Mitzdorf U, Singer W (1978) Prominent excitatory pathways in the cat visual cortex (A 17 and A 18): A current source density analysis of electrically evoked potentials. Experimental Brain Research 33 Available at: http://dx.doi.org/10.1007/bf00235560.

Mitzdorf U, Singer W (1979) Excitatory synaptic ensemble properties in the visual cortex of the macaque monkey: a current source density analysis of electrically evoked potentials. J Comp Neurol 187:71–83.

Muller L, Piantoni G, Koller D, Cash SS, Halgren E, Sejnowski TJ (2016) Rotating waves during human sleep spindles organize global patterns of activity that repeat precisely through the night. Elife 5 Available at: http://dx.doi.org/10.7554/eLife.17267.

Muller L, Reynaud A, Chavane F, Destexhe A (2014) The stimulus-evoked population response in visual cortex of awake monkey is a propagating wave. Nat Commun 5:3675.

Nandy AS, Nassi JJ, Reynolds JH (2017) Laminar Organization of Attentional Modulation in Macaque Visual Area V4. Neuron 93:235–246.

Pereda E, Gamundi A, Rial R, González J (1998) Non-linear behaviour of human EEG: fractal exponent versus correlation dimension in awake and sleep stages. Neurosci Lett 250:91–94.

Plomp G, Michel CM, Quairiaux C (2017) Systematic population spike delays across cortical layers within and between primary sensory areas. Sci Rep 7:15267.

Riera JJ, Ogawa T, Goto T, Sumiyoshi A, Nonaka H, Evans A, Miyakawa H, Kawashima R (2012) Pitfalls in the dipolar model for the neocortical EEG sources. J Neurophysiol 108:956–975.

Schroeder C (1998) A spatiotemporal profile of visual system activation revealed by current source density analysis in the awake macaque. Cerebral Cortex 8:575–592 Available at: http://dx.doi.org/10.1093/cercor/8.7.575.

Schroeder CE, Tenke CE, Givre SJ, Arezzo JC, Vaughan HG Jr (1991) Striate cortical contribution to the surface-recorded pattern-reversal VEP in the alert monkey. Vision Res 31:1143–1157.

Self MW, van Kerkoerle T, Supèr H, Roelfsema PR (2013) Distinct roles of the cortical layers of area V1 in figure-ground segregation. Curr Biol 23:2121–2129.

Senzai Y, Fernandez-Ruiz A, Buzsáki G (2019) Layer-Specific Physiological Features and Interlaminar Interactions in the Primary Visual Cortex of the Mouse. Neuron 101:500–513.e5.

Smith MA, Jia X, Zandvakili A, Kohn A (2013) Laminar dependence of neuronal correlations in visual cortex. J Neurophysiol 109:940–947.

Szymanski FD, Rabinowitz NC, Magri C, Panzeri S, Schnupp JWH (2011) The laminar and temporal structure of stimulus information in the phase of field potentials of auditory cortex. J Neurosci 31:15787–15801.

Takeuchi D, Hirabayashi T, Tamura K, Miyashita Y (2011) Reversal of interlaminar signal between sensory and memory processing in monkey temporal cortex. Science 331:1443–1447.

Wang T, Li Y, Yang G, Dai W, Yang Y, Han C, Wang X, Zhang Y, Xing D (2020) Laminar Subnetworks of Response Suppression in Macaque Primary Visual Cortex. J Neurosci 40:7436–7450.

